# BIOPHYSICAL MECHANISMS UNDERLYING THE GENERATION AND MAINTENANCE OF RULE-LEARNING ENGRAM

**DOI:** 10.1101/2025.08.12.670009

**Authors:** Sankhanava Kundu, Daniel keren, Samma Zidan, Amit Kumar, Jackie Schiller, Edi Barkai

**Author notes:** Corresponding author: Dr. Edi Barkai, Sagol Department of Neurobiology, Faculty of Natural Sciences Haifa University, Haifa 31905, Israel. The last two authors are equal contributors.

## Abstract

Training rodents in a particularly difficult olfactory-discrimination task results with acquisition of high skill to perform the task superbly, termed ‘rule-learning’.

We show that rule-learning occurs abruptly, in a “light bulb moment”. Using whole-cell patch-clamp recordings in the piriform cortex of Fos2A-iCreER/TRAP2 mice, we targeted activated neurons, expressing immediate early genes (IEG).

From the onset of training, IEG-positive neurons from trained animals display enhanced intrinsic excitability. Subsequently, synaptic excitation and inhibition is enhanced in these neurons, in a coordinated, cell-wide process.

In parallel, the density of IEG-expressing neurons sharply declines. Double labeling with TRAP and c-Fos reveals that nearly two-thirds of the rule-memory engram neurons were engaged from the beginning of training. Silencing TRAP-expressing neurons using DREADDs leads to a complete loss of rule memory.

We propose that rule learning occurs at a discrete moment, and is developed through a gradual process that stabilizes the memory of the rule.

## INTRODUCTION

The ability to extract generalizable rules from specific experiences is a fundamental attribute of higher-order cognition. This process, referred to as rule learning or meta-learning enables individuals to “learn how to learn.” In rodents, training on a particularly challenging olfactory discrimination (**OD**) task induces a dramatic acceleration in the learning of novel odorants, suggesting the acquisition of a rule-learning capability (Saar et al. 1998, 1999; Chandra and Barkai 2018; 2019).

A well-established mechanism underlying learning-induced behavioral changes is the modification of intrinsic neuronal excitability. In the piriform cortex, rule learning in OD tasks enhances excitability of pyramidal neurons (Saar et al., 1998; 2001, Chandra and Barkai 2018; 2019). Similar changes have been reported in hippocampal neurons following classical conditioning of the trace eyeblink response (Moyer et al., 1996; Thompson et al., 1996), Morris watermaze task (Oh et al., 2003), and OD learning (Zelcer et al., 2006). Such enhanced excitability is manifested in reduced spike frequency adaptation (Moyer et al., 1996; Thompson et al., 1996; Saar et al., 2001) which is mediated by medium and slow afterhyperpolarizations (AHPs). These AHPs result from potassium currents, which develop following a burst of action potentials (Schwindt et al. 1988; Madison and Nicoll 1984; Constanti and Sim 1987; Saar et al. 2001). In both hippocampal and piriform pyramidal neurons, learning is associated with a reduction in post-burst AHP amplitude, resulting in increased action potential output during sustained depolarization (Moyer et al., 1996; Saar et al., 1998; 2001).

OD rule learning-induced long-term AHP reduction is mediated by reduction in the conductance of the slow calcium-dependent potassium current (sI_AHP_) (Brosh et al. 2006; Saar et al. 2001).

Specifically, the current reduced by OD rule learning is the cholinergic muscarinic current (M-current) (Saar et al. 2001; Awasthi et al. 2022). OD learning-induced AHP reduction in the piriform is also PKC and ERK-dependent (Cohen-Matsliah et al. 2007).

A causal link between rule learning and AHP reduction has been demonstrated by metabotropic activation of the GluK2 Kainate receptor (Chandra et al. 2019); brief activation of GluK2 by direct kainate application or by repetitive synaptic stimulation, results in long-lasting enhancement of neuronal excitability in piriform cortex neurons from controls, but not from OD trained animals (Chandra et al., 2019). Moreover, GluK2 knockout mice show a complete loss of this plasticity and are unable to acquire complex OD tasks that require rule learning, although they retain the ability to perform simple odor discriminations (Chandra et al. 2019). Importantly, viral-induced overexpression of Gluk2 in piriform cortex pyramidal neurons enhances rule learning remarkably. Unlike synaptic modifications, which are input-specific, a change in the neuron’s firing patterns globally affect the neuron. This broad enhancement of excitability are expected to increase the amount of information (expressed in the average action potential firing rate) processed in the network. In particular, the enhanced intrinsic neuronal excitability results in enhanced learning abilities (Lisman et al., 2018, Chandra et al, 2019). Such a state, characterized by heightened excitability, may functionally correspond to a “*learning mode*,” during which neurons are more easily recruited into memory traces through activity-dependent synaptic modifications (Yiu et al., 2014; Gouty-Colomer et al., 2016; Chandra et al., 2019; Lisman et al., 2018). Following OD rule learning, this high-excitability state persists in the piriform cortex for several days, correlating with enhanced performance on tasks requiring rule acquisition (Saar et al. 1998; 2001 Chandra and Barkai 2018).

Despite this knowledge, it remains unclear how rule learning–induced increases in intrinsic excitability are accompanied by changes in synaptic connectivity within each neurons and what is the timeline of these changes. Evidence supporting such a combined mechanism comes from recent studies showing that subpopulations of neurons with high baseline firing rates displayed both increased intrinsic excitability and recurrent synaptic connectivity (Trojanowski et al., 2021). In the context of OD learning, rule leaning is accompanied by enhanced synaptic excitation and inhibition onto pyramidal neurons, and increase in spine density, both in recurrent synapses at the proximal apical dendrite of the piriform cortex neurons and in CA1 pyramidal neurons (Saar et al. 2012, Ghosh et al., 2015; 2016, Knafo et al. 2004; 2005a; 2005b).

Here we combine whole cell patch clamp recordings with identification of active neurons using immediate early genes (IEGS) expression to describe the cellular and network biophysical mechanisms underlying acquisition of rule learning and its long-term memory maintenance.

We first show that rule learning occurs at within a sharply defined temporal window, which may be best termed as “light-bulb moment”. We also show that the rule learning engram is supported by a subgroup of neurons, most of which were activated early during learning. These neurons are initially intrinsically more excitable and subsequently develop enhanced synaptic connectivity, both excitatory and inhibitory. This coordinated sequence of intrinsic and synaptic modifications enables the formation and stabilization of the long-term memory engram for the rule.

## RESULTS

### Rule learning is achieved after prolonged training period and becomes apparent in a well-identified “light bulb moment”

We trained mice for OD discrimination rule in a 4-arm maze (Figure 1A). When examining the group performance across training sessions (n=63 mice) we observed a gradual improvement culminating in all mice successfully acquiring the rule (Figure 1B). However, individual learning rates varied considerably, ranging from 5 to 11 days—with two outliers requiring 13 and 14 days, respectively—and an overall average of 8.4 ± 1.8 days (Figure 1C).

**Figure 1:**
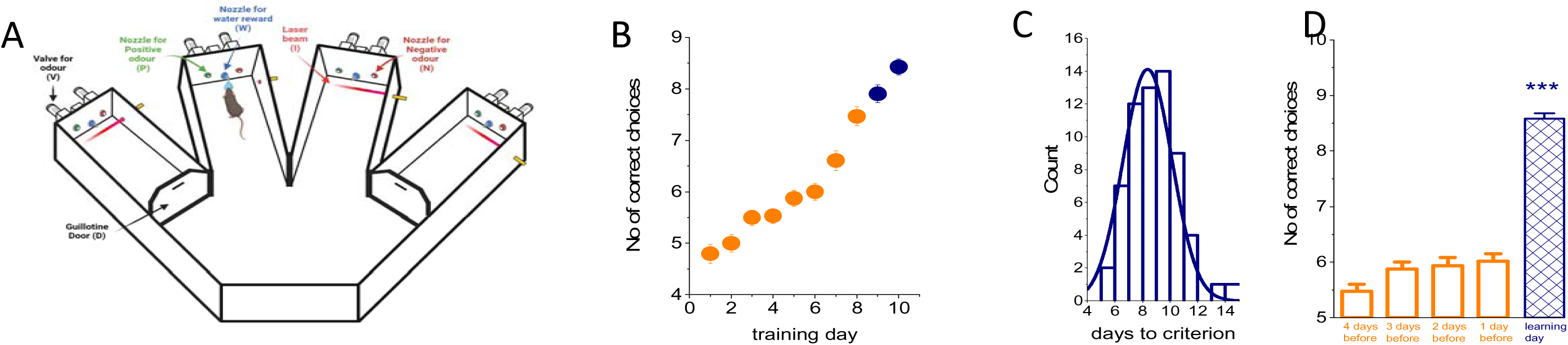
Rule learning in the complex OD task appears to occur in a step function. **A.** Schematic description of the 4-arm maze. V=odor and water valves, P= positive-cue odor, N= negative-cue odor, D= doors, I= infrared beam, W= water reward. **B.** A curve describing the averaged success rate of all 63 mice in the course of training for the rule. Values in **navy** denote the day in which the averaged performance met the criterion for rule leaning. **C.** A normal distribution curve showing the number of days required to achieve the criterion for rule learning. The averaged time was 8.4±1.8 days. **D.** Columns show the average improvement in correct choices made by the 63 mice on the final day, when the rule was acquired (column in **navy**) and during the four previous training days (**orange**).Values represent mean ± SE. (*** P<0.001)

To account for this variability, we aligned training performance for each mouse to its own final day of training and analyzed progression relative to that point (Figure 1D).

Our data show that during the four days preceding rule acquisition, mice improved their performance only marginally (from 5.5±1.0 correct choices on the fourth day before learning to 6.0±1.0 on the last day before learning). In contrast, on the final training day—when each animal reached the rule-learning criterion, mice improved the number of their correct choice by an average to 8.6±0.7 correct choices. Thus, the improvement on the last day was more than 6.5 times higher than the cumulative improvement observed over the prior four days, indicating the occurrence of a distinct “light bulb moment”, when the animal abruptly grasps how to perform the task successfully. Notably, we did not find any difference in performance between females and males. Therefore, data obtained using the two genders were combined.

### Learning-induced AHP reduction cannot be explained by a whole network effect on a normally distributed neuronal population

Previous studies showed that learning, in particular complex learning that requires prolonged training periods, results in a whole network reduction of the post-burst AHP in the pyramidal cell population. This was shown in piriform cortex neurons following operant conditioning (Saar et al., 1998; 2001, Saar and Barkai 2003) (Figure 2A). This effect is also apparent in hippocampal neurons following classical conditioning of the trace eyeblink response (Moyer et al., 1996; Thompson et al., 1996), following training in the water maze task (Oh et al. 2003), and following OD learning (Zelcer et al. 2006). However, previous analysis leading to the interpretation of the “whole network effect” assumed that the post burst AHP is distributed normally in the pyramidal cell population in the piriform cortex (Saar et al., 1998; 2001) and the hippocampus (Moyer et al. 1996). We reexamined this assumption by performing a meta-analysis of AHP amplitudes reported in our previous studies (Saar et al. 1998; 2001, Seroussi et al. 2002, Brosh et al. 2006; 2007, Cohen-Matsliah et al. 2007; 2009, Chandra et al. 2019, Awasthi et al. 2022) added to additional neurons recorded recently. Altogether, we collected 128 neurons from naïve rats, 132 neurons from trained rats (recorded three days after rule learning) and 137 neurons from pseudo-trained rats.

**Figure 2:**
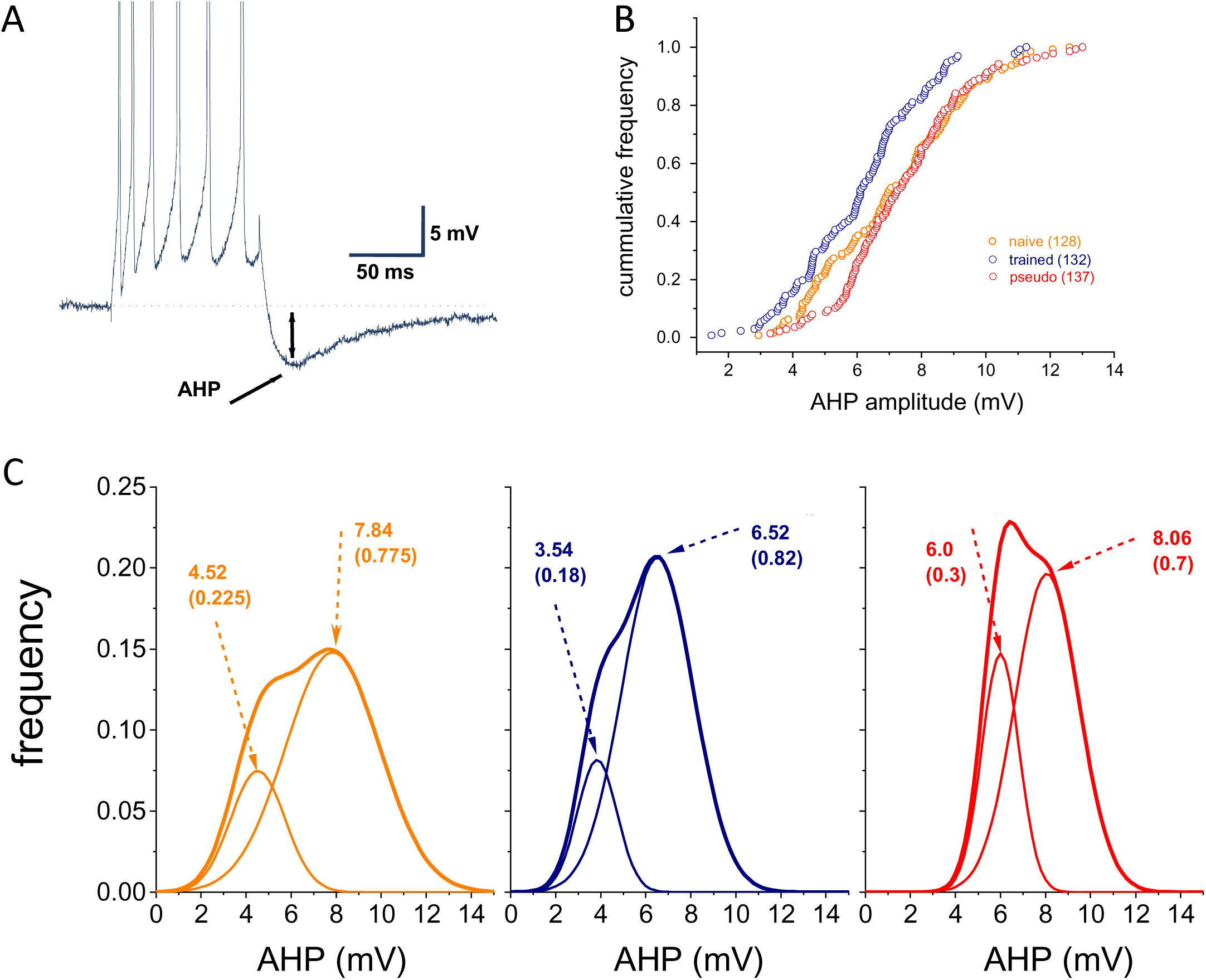
The post burst AHP distribution in neurons from trained mice entails two populations. **A.** Current clamp recordings with sharp electrodes for AHP measurements in a piriform cortex neuron. The holding membrane potential is –60 mV and the AHP is generated by 100 ms depolarizing current step, with intensity that generates a train of 6 action potentials. **B.** Cumulative frequency distributions of the AHP amplitudes in neurons from the three groups. The AHP amplitudes from trained rats curve is smoothly shifted to the left along the x-axis relative to those describing the distributions of AHP amplitudes in neurons from the two control groups, indicating that in most recorded neurons from the trained rats the amplitude of the post-burst AHP is reduced. Data taken from Saar et al. 1998; 2001, Seroussi et al. 2002, Brosh et al. 2006, Cohen-Matsliah et al. 2007, Chandra et al. 2019, Awashti et al. 2022. **C.** Distributions of the post burst AHP’s amplitudes for the three groups. For each group, the best fit was achieved by the combination of two normal distributions. The respective numbers denote the average of each normal distributions and its weight in the population. The difference in the average amplitudes of the two distributions in the trained group (54.3%) is almost identical to the difference in the naïve group (57.7%). Also, the ratio (trained/naive) between the first two peaks (0.78) is similar to the ratio between the second two peaks (0.83). Note that the two peaks of the distributions for the trained group are lower than the parallel peaks in the control groups. These results imply that in most, if not all, neurons in the trained group the AHP was reduced by rule learning. The thick line in each graph describe the sum of the two distributions.

The cumulative frequency distributions shown in figure 2B suggest that there’s a general reduction in the post-burst AHP values in neurons from the trained group compared with the two control groups, and that the values of the pseudo trained and naïve groups are overlapping, except for the neurons with low AHPs.

However, a normality test applied to these three groups showed that the naïve and pseudo trained groups are not distributed normally (p values; 0.035 for the naïve, 0.29 for the trained and 0.025 for the pseudo trained). Thus, we used the Kolmogorov-Smirnov test to determine if the three groups differ from each other. The trained group differs significantly from both the naïve (D=0.2415, p=0.001) and the pseudo trained (D=0.2809, p<0.001) groups. Notably the two control groups also differed (D=0.1858, p=0.0207).

We next examined if the AHP values can be fitted by distribution curves of two Gaussians. The result of this analysis is shown in figure 2C. Our analysis shows that the naïve group is composed of two neuronal populations, one that entails 22.5% of the group with an average AHP value of 4.52 mV and a second group with the remaining 77.5% neurons with an average value of 7.84 mV. The trained group is composed of one population that entails 18% of the group with an average AHP value of 3.54 mV and a second group with the remaining 82% neurons with an average value of 6.52 mV. The pseudo trained group is composed of one population that entails 30% of the group with an average AHP value of 6.0 mV and a second group with the remaining 70% neurons with an average value of 8.06 mV. Moreover, these results suggest that the AHP was reduced in most, if all, neurons in, the trained group (Figure 2C legend).

It has been suggested that neurons are recruited to a memory trace based on relative neuronal excitability immediately before training (Yiu et al. 2014, Gouty-Colomer et al., 2016). Notably in the piriform cortex the proportion of neurons with reduced AHP is similar in the naive and trained groups (Figure 2C). We thus hypothesized that as shown for simpler forms of learning, the engram for long-term memory for rule-learning is also composed from piriform cortex neurons which were more excitable when the long training period in the OD task began, although many training days and sessions would be required for the memory to occur.

### Immediate early genes (IEGS) expressing neurons show enhanced intrinsic excitability and reduced AHP at the onset of training in neurons from trained mice only

Over the past decade, it was demonstrated that learning is associated with the induction of immediate early genes (IEGs), such as cFos and activity-regulated cytoskeleton-associated protein (Arc) (Josselyn et al. 2015; Ryan et al. 2015; Meissner-Bernard et al. 2019). Notably, Gouty-Colomer et al. (2016) demonstrated that in naïve mice, Arc-expressing neurons exhibit significantly higher intrinsic excitability than non-expressing neurons, making them more likely to be recruited into a fear memory trace.

We used an advanced version of targeted recombination in active populations (TRAP2) mouse line, in which c-Fos expression drives tamoxifen-inducible Cre recombination to obtain genetic access to neurons that were activated during rule-learning (Guenthner et al. 2013) (Figure 3). In layer II pyramidal neurons of the piriform cortex, we found no significant difference in intrinsic excitability between tagged and non-tagged neurons in naïve mice. Tagged neurons fired 18.8 ± 3.0 spikes (n = 17) in response to somatic current steps, compared to 18.6 ± 4.9 spikes (n = 18) in non-tagged neurons (t = 0.19, p = 0.8) (Figure 4A, B).

**Figure 3:**
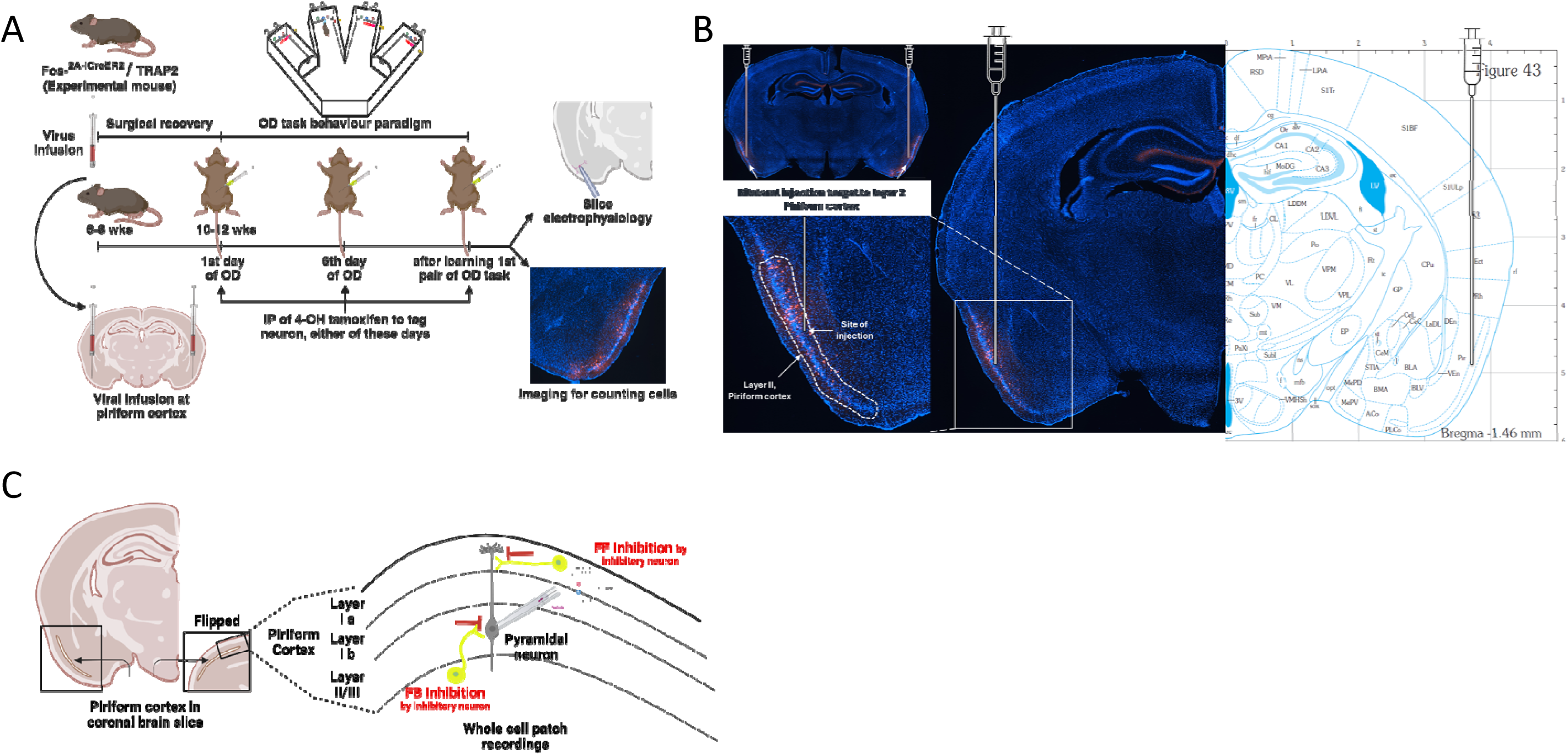
Schematic description of the overall experimental procedure, viral infection site and whole-cell patch recording site. **A.** General approach to mark the activity dependent cell marking as training progresses and rule learning is maintained, using TRAP2 mice strain. Six-eight weeks old TRAP2 mice were used for virus injection (AAV-8/2-hSyn1-dlox-tdTomato(rev)-dlox-WPRE-bGHp(A); VVF-Zurich) in layer II Pyramidal neurons. After recovering from the surgery and time allowed for the expression of the virus, mice were trained in the olfactory maze. On the chosen day, before or during rule learning, tamoxifen (IP; 5mg/KG body weight) were injected to allow the expression of Cre recombinase and the expression of reporter to tag the cells. Following that, animals were used as per experimental need. **B.** The Injection site in layer-II piriform cortex shown in one brain hemisphere, compared with the other hemisphere from “Mouse brain atlas” (Franklin & Paxinos; 3^rd^ Edition) with a Closeup view of the injection site. Active cells are expressed in red (tdTom), within layer II piriform cortex **C.** Schematic illustration of the whole cell recordings (a recording electrode (grey) used to place in piriform layer II pyramidal cell bodies for various recordings).

**Figure 4:**
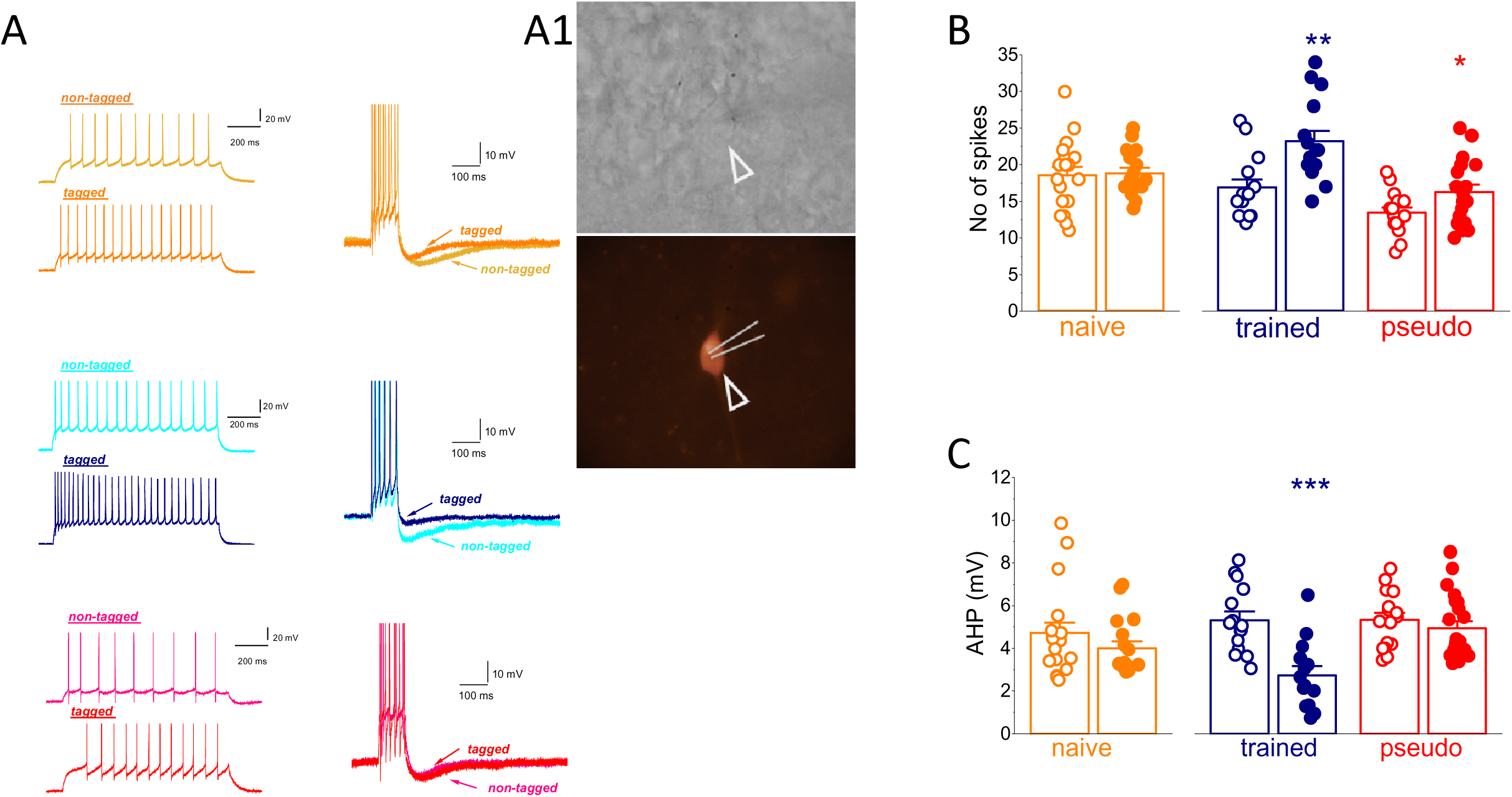
neurons expressing IEG after one day of training show enhanced intrinsic excitability and reduced AHP. **A.** Examples of repetitive spike firing (left) and AHP amplitudes (right) after the first day of training in the three groups. For each group, examples of recorded traces are shown from non-tagged and tagged neurons. A1: representative images of whole-cell patch of tagged neuron under DIC and green filter. **B.** Tagged neurons have enhanced neuronal excitability in both groups (trained and pseudo trained) that were exposed to the maze, regardless if mice were trained for rule learning. **C.** Tagged neurons from trained mice only show reduced AHP compared to their non-tagged neighbors. This AHP reduction in IEGs expressing neurons is related to training for the rule. Each point represents the average number of spikes (B) or the AHP amplitude (C) for each neuron. Bars represent mean ± SE. (**, P<0.01; *** P<0.001).

In contrast, in trained mice that had completed just the first day of training, tagged neurons fired significantly more action potentials (23.2 ± 5.6 spikes; n = 15) than non-tagged neurons (16.9 ± 4.0 spikes; n = 15) (t = 3.49, p = 0.016). Similarly, in pseudo-trained mice—exposed to the training context but not the learning contingency—tagged neurons were also more excitable (16.3 ± 4.4 spikes; n = 19) than non-tagged neurons (13.4 ± 2.9 spikes; n = 16) (t = 3.49, p = 0.016). Importantly, the number of action potentials in tagged neurons from trained mice was significantly higher than in neurons from both naïve (t=2.77, p=0.009) and pseudo trained mice (t=4.05, p=0.003). The number of action potentials in the naïve group was almost significantly higher than in the pseudo trained group ((t=2.0, p=0.053).

Thus, although the first day of training does not result in improved performance in the OD maze (see figure 1B), tagged neurons show enhanced intrinsic excitability in the trained group, particularly as do, to a lesser extent, neurons for pseudo-trained mice (Figure 4B). These data suggest that the neurons recruited for learning are not those that already exhibited higher excitability prior to training. Enhancement of intrinsic excitability appears to occur in these neurons only in the context of actual learning. Interestingly, the OD pseudo training procedure appears to have the opposite effect of reducing the intrinsic excitability of piriform cortex pyramidal neurons.

Such enhanced intrinsic excitability was accompanied by reduction in the post burst AHP in tagged neurons from trained mice only (Figure 4A, C). In naïve mice, AHP amplitude did not differ significantly between tagged (4.0 ± 1.3 mV; n = 17) and non-tagged neurons (4.7 ± 2.1 mV; n = 18) (t = 1.16, p = 0.25). In contrast, in trained mice, tagged neurons showed a significantly reduced AHP (2.8 ± 1.6 mV; n = 14) compared to non-tagged neurons (5.3 ± 1.6 mV; n = 15) (t = 4.4, p = 0.00015). No significant difference in AHP was observed between tagged and non-tagged neurons in pseudo-trained mice (4.9 ± 1.5 mV vs. 5.3 ± 1.3 mV; t = 0.79, p = 0.43).

These findings show that: ***a.*** the averaged amplitude of the AHP in tagged neurons from trained mice is 52% compared with non-tagged neurons. This difference is similar to the one observed between the two normally distributed groups of neurons shown for naïve animals (58%, see legend for figure 2C). This suggests that neurons recruited for rule learning are those with initially lower AHPs, consistent with previously work in fear conditioning (Gouty-Colomer et al., 2016). ***b.*** the AHP amplitude of the tagged neurons in trained mice is significantly lower than that of tagged neurons in naïve mice (t=2.4, p=0.023). Thus, once recruited for rule learning, neurons undergo further enhancement of their intrinsic excitability.

### Enhanced intrinsic excitability and reduced AHP in IEGS expressing neurons from trained mice persists throughout the training period

We next examined if such enhanced excitability and reduced AHP is apparent in IEGs expressing neurons throughout the prolonged training period. We measured the No of spikes and AHP values in tagged and non-tagged neurons on the sixth day during ongoing training and after the completion of rule learning. Our results show that enhanced excitability in tagged neurons from trained mice persists during rule learning (Figure 5A_1_); tagged neurons from trained mice that completed the six days of training fired a significantly higher number of spikes compared with non-tagged neurons [21.7±5.5 (n=21) spikes for tagged neurons compared with 17.9±3.7 (n=20) for non-tagged neurons, t=2.57, p=0.014]. Also, tagged neurons from pseudo-trained mice that completed the six days of training fired a significantly higher number of spikes compared with non-tagged neurons [20.1±3.2 (n=12) spikes for tagged neurons compared with 16.8+4.8 (n=16) for non-tagged neurons, t=2.7, p=0.011].

**Figure 5:**
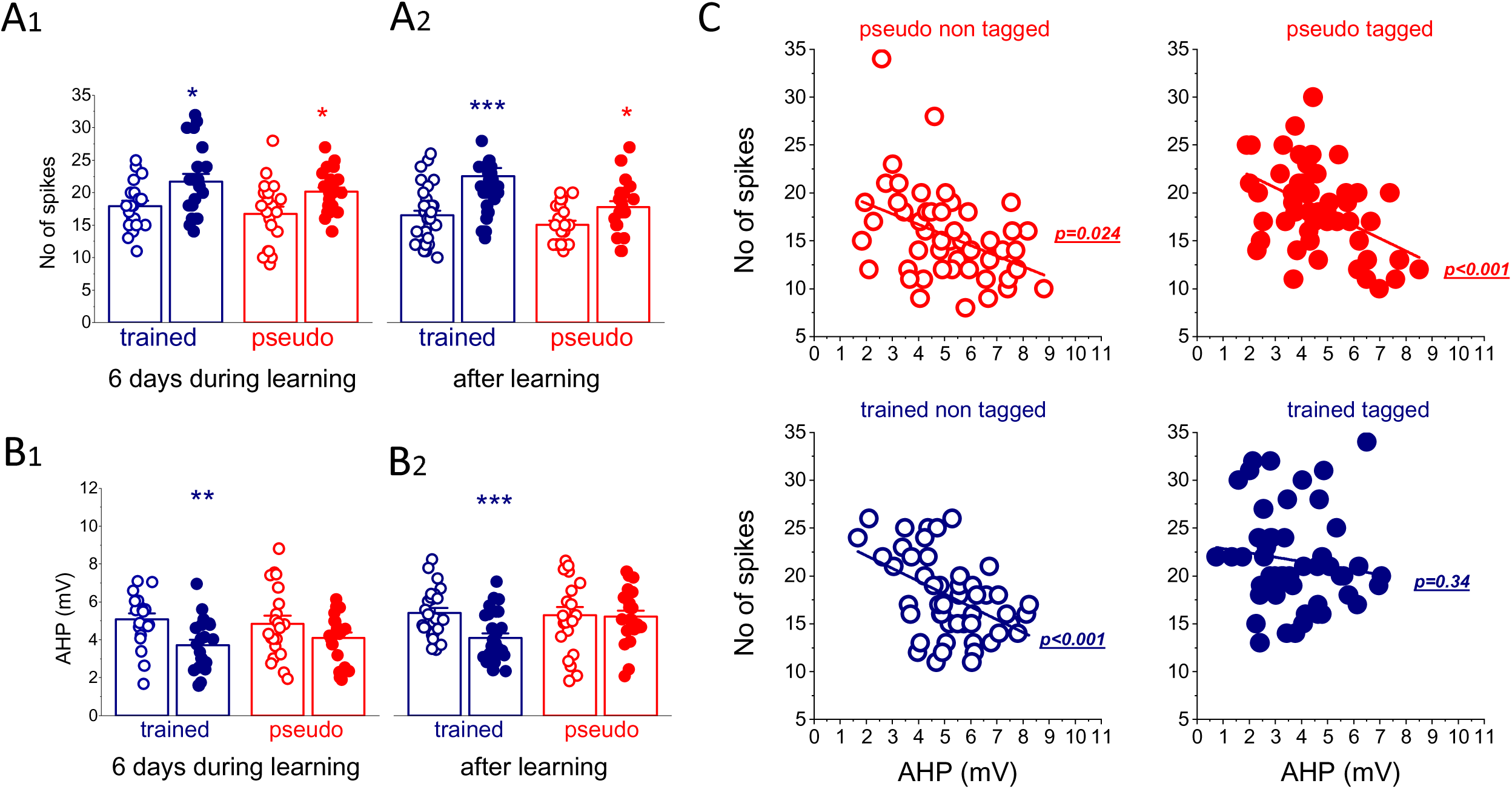
Enhanced intrinsic excitability in trained and pseudo trained and reduced AHP in trained in tagged neurons persists throughout the training period. **A.** Number of spikes generated by tagged and non-tagged neurons from the three groups six days after the beginning of training (A_1_) and after rule learning (A_2_). Enhanced neuronal excitability in tagged neurons is evident throughout the training period in both groups that were exposed to the OD maze. **B.** The average AHP amplitude is reduced in neurons from trained mice only. This effect also persists from the beginning of training and remains after rule learning is completed. **C.** A clear correlation exists between the AHP amplitudes in neurons from all groups, except for the tagged neurons in the trained group. This result implies that in tagged trained neurons the AHP is reduced to the point where it no longer controls intrinsically evoked respective firing. Each point represents the average number of spikes (A) or the AHP amplitude (B) for each neuron. Bars represent mean ± SE. (*, P<0.05; **, P<0.01; *** P<0.001).

After the completion of learning (one to three days after rule learning is achieved), tagged neurons from trained mice fired a significantly higher number of spikes compared with non-tagged neurons [22.5±7.5 (n=35) spikes for tagged neurons compared with 16.5±4.3 (n=33) for non-tagged neurons, t=4.04, p=0.00014] (Figure 5A_2_). Also, tagged neurons from pseudo-trained mice fired a significantly higher number of spikes compared with non-tagged neurons [17.8±4.2 (n=20) spikes for tagged neurons compared with 15.0±3.0 (n=12) for non-tagged neurons, t=2.4, p=0.023]. Notably, tagged neurons from trained mice had a significantly higher number of spikes, compared with tagged neurons from pseudo trained mice (t=2.61, p=0.003). No such difference was found for non-tagged neurons between the two groups (t=1.47, p=0.015).

Here too, enhanced intrinsic excitability was accompanied by reduction in the post burst AHP in tagged neurons from trained mice only (Figure 5B).

After six days of learning, tagged neurons from trained mice had a significantly lower AHP (3.7±1.4 mV (n=21)), compared with 5.1±1.4 mV (n=20) from non-tagged neurons (t=3.2, p=0.003) (Figure 5B1). In tagged neurons from pseudo trained mice the amplitude of the post-burst AHP was 4.1±1.3 mV (n=21), compared with 4.9±1.3 (n=21) for non-tagged neurons (t=1.54, p=0.13). After rule leaning completion, tagged neurons from trained mice had a significantly lower AHP (4.1±1.3 mV (n=29), compared with 5.4±1.3 mV (n=25) for non-tagged neurons (t=3.28, p=0.0004) (Figure 5B). In tagged neurons from pseudo trained mice the amplitude of the post-burst AHP was 5.0±2.0 mV (n=10), compared with 5.2±2.2 (n=14 for non-tagged neurons (t=0.25, p=0.8).

Thus, throughout the prolonged training period, IEGs expressing neurons in trained mice showed enhanced excitability and reduced AHP. Additionally, neurons from pseudo trained mice also showed enhanced excitability, but to a significantly lesser extent and without any significant changes in the AHP amplitude.

As shown in figure 5C, the AHP reduction in tagged neurons from trained mice results in the loss of its ability to control repetitive firing in neurons, which may account for the enhanced excitability in these neurons compared with both IEGs expressing neurons in pseudo trained mice and IEGs non-expressing neurons in the PC of trained mice.

### Learning-induced synaptic enhancement reflects selective recruitment, not a global network shift

Similarly to what was shown for intrinsic excitability, previous studies showed that OD complex learning results in global enhancement of both synaptic inhibition and excitation onto the piriform cortex pyramidal neurons (Saar et al. 2012, Ghosh et al., 2015; Brosh et al. 2009, Kfir et al, 2014, Kundu et al. 2024, Reuveni et al., 2013; 2017) (Figure 6A-B). Although most neurons exhibited increased postsynaptic current amplitudes, a subset showed disproportionately large increases in miniature event amplitudes (Saar et al., 2002; Ghosh et al., 2015, 2016) (Figure 6B), suggesting non-uniform changes across the population.

**Figure 6:**
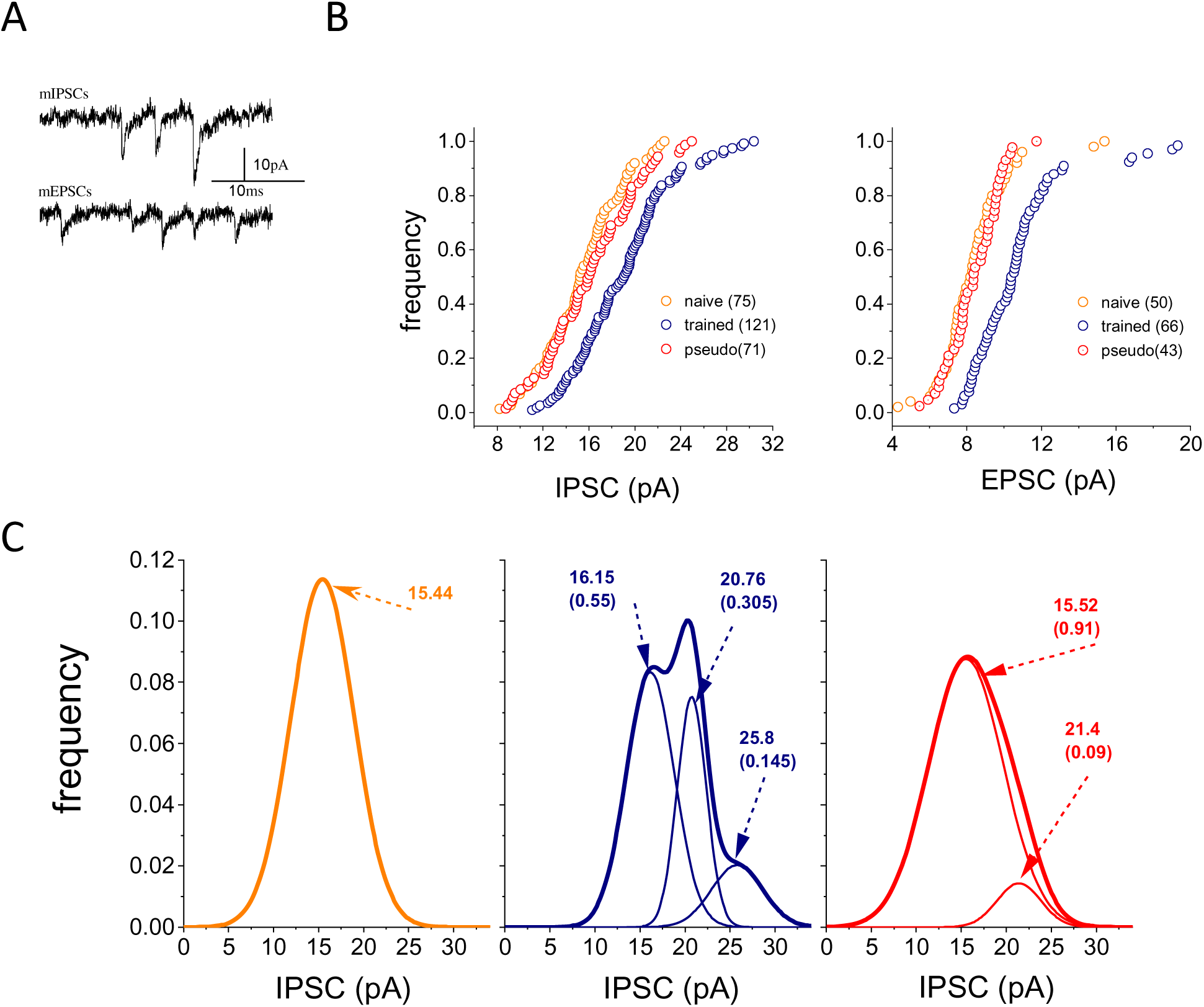
OD rule learning-induced enhancement of synaptic inputs onto PC pyramidal neurons is not uniform. **A.** Examples of recordings of mIPSCs and mEPSCs from PC pyramidal neurons (see Ghosh et al. 2015, 2016 for details of recordings conditions) **B.** Cumulative frequency distributions of the IPCSs (left) and the EPSCs (right) amplitudes in neurons from the three groups. The amplitude curves of both inhibitory and excitatory synaptic events from trained rats are smoothly shifted to the right along the x-axis relative to those describing the distributions of synaptic currents in neurons from the two control groups, indicating that in most recorded neurons from the trained rats the amplitude of both synaptic inhibition and excitation are enhanced. Data taken from Saar et al. 2002, Ghosh et al. 2015; 2016; 2018, Jammal et al. 2018, Kundu et al. 2024. **C.** Distributions of the mIPCSs amplitudes for the three groups. For the naive group, the best fit was achieved by a single normal distribution. Values for the trained group could be fitted by three normal distributions. Values for the pseudo trained group could be fitted by two normal distributions, although most neurons (91%) belong to the first distribution. The respective numbers denote the average of each normal distributions and its weight in the population. Notably, the peaks of the first (lowest) distributions (the single distribution for the naive group) are almost identical. Thus, all rule learning-induced enhancements of synaptic inhibition target 45% of the pyramidal neurons that compose the second and third distributions.

We thus examined if the strength of synaptic inhibition is distributed normally. Here, the normality test applied to the three groups showed that the naïve and pseudo trained groups are distributed normally (p values; 0.66 for the naïve and 0.44 for the pseudo trained). The trained group is not distributed normally (p=0.03). Thus, we used the Kolmogorov-Smirnov test to determine if the three groups differ from each other. The trained group differs significantly from both the naïve (D=0.384, p=3×10^-6^) and the pseudo trained (D=0.2728, p=0.0001) groups, whereas the two control groups did not differ (D=0.1319, p=0.3658).

Normality test applied for synaptic excitation showed that the naïve group (p=3.7×10^-4^) and the trained group (p=3.8×10^-8^) are not distributed normally. The pseudo trained group shows normal distribution (p=0.87). The Kolmogorov-Smirnov test shows that the trained group differ significantly for the naïve (D=0.497, P<0.001) and the pseudo trained groups (D=0.521, P<0.001) groups. Here too, the two control groups did not differ (D=0.1274, p=0.847).

To better characterize the synaptic distributions, we fitted IPSC amplitudes (averaged per neuron) to Gaussian mixture models using recordings five days post-training (Saar et al. 2012). The naïve group (n = 75) was best described by a single population (mean = 15.44 pA). In contrast, the trained group (n = 123) was composed of three distinct populations: 55% of neurons with a mean IPSC of 16.15 pA, 30.5% with 20.76 pA, and 14.5% with 25.8 pA. The pseudo-trained group (n = 71) included one major population (91% of neurons, mean = 15.52 pA) and a minor subpopulation (9%, mean = 21.4 pA) (Figure 6C). Thus, it may be concluded that 45% of the PC neurons show enhanced inhibitory inputs five days after rule learning.

### Synaptic transmission onto tagged neurons only is enhanced after rule learning

We next examined whether tagged neurons have stronger synaptic inputs. We compared the averaged amplitudes of mEPSCs and mIPCSs in tagged and non-tagged neurons from the three groups. Neurons were recorded with cesium methanesulfonate containing electrodes (see methods). Each neuron was voltage clamped at two different holding potentials: at –70 mV to record mEPSCs and at 0 mV to record mIPSCs as previously described (Kundu et al. 2024) (Figure 7A).

**Figure 7:**
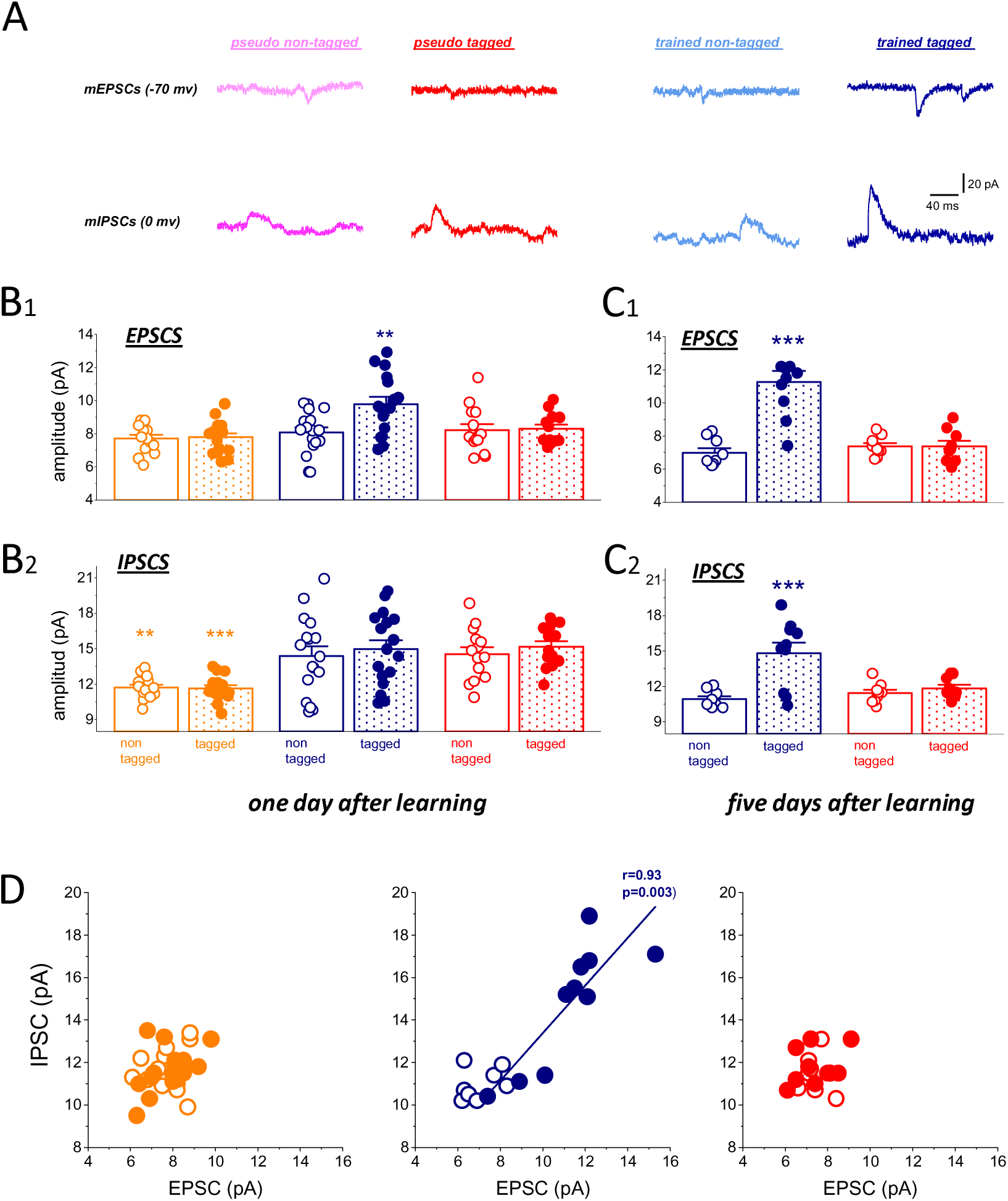
rule learning induced enhancement of excitatory and inhibitory synaptic transmission is positively correlated in trained tagged neuron. **A.** Examples of a spontaneous inhibitory and an excitatory synaptic event recorded from the same neurons **B.** Miniature excitatory (B_1_) and inhibitory (B_2_) synaptic events amplitude one day after rule learning completion. At this point in time the synaptic excitation was enhanced in IEGs expressing neurons from trained mice only. Synaptic inhibition was enhanced in all neurons from mice that were exposed to the maze and odors, regardless if they were trained to learn the rule. Each point represents the average synaptic event amplitude for each neuron. Bars represent mean ± SE. (**, P<0.01). **C.** Miniature excitatory (C_1_) and inhibitory (C_2_) synaptic events amplitude five day after rule learning completion. At this point in time both synaptic excitation and inhibition were enhanced for IEGs expressing neurons from trained mice only In **B**&**C**, Each point represents the averaged synaptic event amplitude for each neuron. Bars represent mean ± SE. (***, P<0.001). **D.** Five days after rule learning, a positive strong correlation between the averaged mEPSCs and mIPSCs in each cell in IEG-expressing neurons from trained mice only (p values for linear fit were 0.63 for naïve non-tagged, 0.09 for naïve tagged, 0.31 for trained non-tagged, 0.94 for pseudo trained non-tagged and 0.34 for pseudo trained tagged neurons).

One day after rule learning, learning-induced enhancement of synaptic excitation was apparent in these recording conditions in tagged neurons from trained mice only (Figure 7B_1_). The averaged mEPSC amplitude in these neurons (9.8±1.8 pA, n=17 neurons) was significantly (F_2,45_=10.1, p<0.001) higher compared to the averaged value in neurons from pseudo-trained (8.3±0.9 pA, n=14) and naïve mice (7.8±1.0 pA, n=16). Post hoc t-tests show that the value for trained neurons was significantly higher compared with both naïve (p<0.001) and pseudo trained (p=0.009) neurons. The average values of the two control groups did not differ (p=0.14). Also, the averaged value of trained tagged neurons was significantly higher than in non-tagged neurons (8.0±1.3 pA, n=15) from the same group (p=0.0083). No difference was found between non-tagged neurons from the three groups.

At the same point in time (one day after training completion), the amplitudes of the mIPSC from naïve neurons were smaller compared to the neurons from trained and pseudo trained mice (Figure 7B_2_). This was apparent for both non-tagged (F_2,45_=12.7, p=0.000046) and tagged neurons. Thus, training in the complex OD task results in a temporary increase in synaptic inhibition which occurs throughout the pyramidal cell population (tagged and non-tagged). Likely, such increase is related to the exposure to the maze and odors, rather than to the rule learning.

In contrast, five days after rule learning, the point in time when the enhancement of learning-induced synaptic transmission is at its peak (Saar et al., 2012, Kundu et al. 2024, Reuveni et al., 2013; 2017), learning-induced enhancement of both synaptic excitation and inhibition were apparent in tagged neuron from trained mice only (Figure 7C_1_, C_2_).

The averaged mEPSC amplitude in these neurons (11.2±2.1 pA, n=10 neurons) was significantly (F_2,33_=23.9, p<0.001) higher compared to the averaged value in neurons from pseudo-trained (7.4±1.0 pA, n=9) and naïve mice. Post hoc t-tests show that the value for trained neurons was significantly higher compared with naïve (p<0.001) and pseudo trained (p<0.001) neurons. Also, the averaged value of trained tagged neurons was significantly higher compared to non-tagged neurons (7.0±10.8 pA, n=9) from the same group (p<0.001). No difference was found between non-tagged neurons from the three groups.

Also, the averaged mIPSC amplitude from the trained tagged neurons (14.8±2.9 pA, n=10 neurons) was significantly (F_2,33_=11.2, p<0.001) higher compared to the averaged value in neurons from pseudo-trained (11.8±0.9 pA, n=9) and from naïve mice. Post hoc t-tests showed that the value for trained neurons was significantly higher compared with naïve (p<0.001) and pseudo trained (p<0.008) neurons. The averaged value of trained tagged neurons was significantly higher compared with non-tagged neurons (10.9±0.7 pA, n=9) from the same group (p<0.001). No difference was found between non-tagged neurons from the three groups.

As shown in figure 7D, learning induced enhancement of both inhibitory and excitatory synaptic inputs is strongly correlated only in tagged neurons. This finding supports the notion that rule learning drives coordinated changes in synaptic connectivity at the level of individual neurons, rather than through independent modulation of synapses across the network (Saar et al, 2012, Ghosh et al. 2015; 2016, Reuveni et al. 2017).

The convergence of changes in all three key components of neuronal excitability—intrinsic excitability, excitatory synaptic input, and inhibitory synaptic input—highlights these tagged neurons as primary candidates for forming the cellular substrate of the rule-learning engram.

### Dynamics of recruitment and discharge of activated pyramidal neurons during the course of rule learning

We next explored the dynamics of IEGs-tagged neuronal labeling during the course of rule learning. The experimental design is described in the Figure 8A_1_.

**Figure 8:**
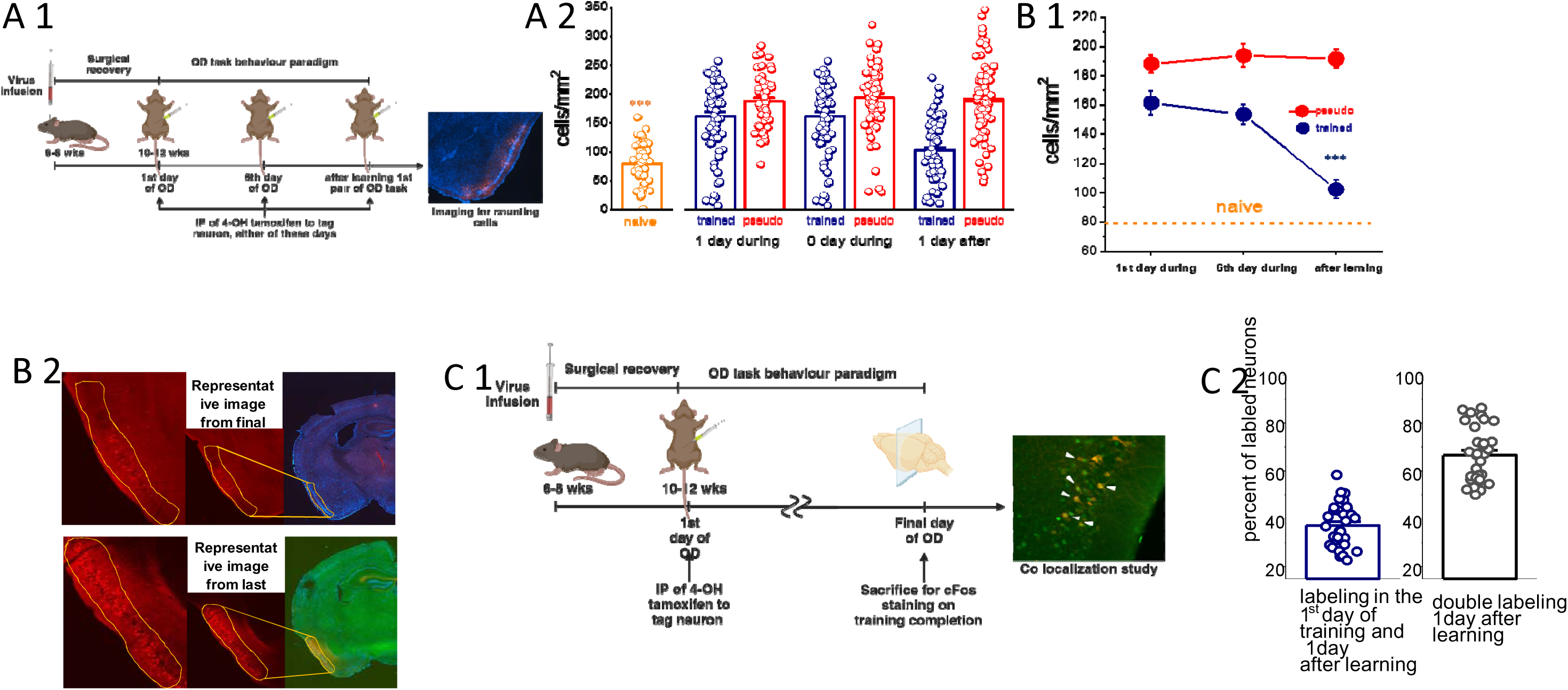
the number of IEGS expressing neurons drops markedly after rule learning completion. **A_1_.** Schematics of experimental approach and division of experimental TRAP animals and procedure to mark the activity dependent neurons by applying 4 OH tamoxifen injection. **A_2_, B_1_.** The density of IEGs-expressing neurons in brains of trained and pseudo trained mice increases at least twofold from the first day of training and throughout the training period. The density of tagged neurons decreases dramatically in brains of trained mice once rule learning is completed. For columns (A2) each point represents the number of cells counted in each slice (A2). Bars represent mean ± SE. The same data is presented in **B1** as averaged values for all slices ± SE (*** P<0.001). **B_2_.** <u>Top</u>: Representative image from final day of training from trained group animal, showing expression of active cell population in the piriform cortex. Yellow line drawn to demark the layer II piriform cortex. <u>Bottom</u>: Representative image from final day of training from pseudo trained group animal, showing expression of active cell population in piriform cortex. Yellow line drawn to demark the layer II piriform cortex. **C.** Dynamics of recruitment and discharge of neurons from activation in the course of rule learning **C_1_.** Schematics of experimental approach for co-localization study. One-month old TRAP mice were used for virus injection (AAV-8/2-hSyn1-dlox-tdTomato(rev)-dlox-WPRE-bGHp(A); VVF-Zurich) in layer II Pyramidal neurons. After recovering from the surgery and time allowed for expression of the virus, mice were trained in the olfactory maze. On the chosen the day, before or during rule learning, tamoxifen (IP; 50mg/KG body weight) were injected to allow the expression of Cre recombinase and expression of reporter to tag the cells. The same animals were chosen to continue the OD training. Later, during or after OD training, the mice were sacrificed 90 minutes after the last training session and perfused to do IHC for c-Fos. c-Fos is mostly expressed in excitatory cells, as these neurons compose the majority of cells in layer II of the piriform cortex (more than 90%, see Suzuki and Bekkers, 2010). The overlay of the marked neurons were detected by fluorescent microscopy. ***picture:*** Double labeling, on the first day of trained and after rule learning, shows that about third of the neurons activated at the very beginning of training remain activated after rule earning is completed (left). **C_2_.** Bar graph showing the cell density (cells/mm^2^.) counted on final day of training by IHC performed for cFOS (green) positive cells along with TRAP positive (red) cells on final day of training from trained group particularly. Each circle represents one hemisphere from one mouse. Two thirds of the neurons that remain active after rule learning were recruited to engram since day one (right). Bars represent mean ± SE.

In naïve animals, the density of tagged neurons was 80.8±33.3 cells/mm^2^ (n=44 slices; Figure 8A_2_). This was significantly lower (F_2,105_=50.4, p<0.001) than the density observed one day after training began in both pseudo-trained (188.1 ± 45.1 cells/mm², n = 53) and trained mice (161.4 ± 68.8 cells/mm²). Interestingly, pseudo-trained animals had a significantly higher density of tagged neurons than trained animals (t = 2.45, p = 0.0103). These results indicate that mere exposure to the maze and odors, independent of learning, induces approximately a twofold increase in the number of IEG-tagged neurons.

To explore how the number of labeled neurons is modified during the course of rule learning, we compared the pseudo trained and the trained groups at two additional time points, at the sixth day of learning and one day after the mice learned the rule (experimental procedure figure 8A_1_). The number of tagged cells in pseudo trained brains remained stable, almost identical, for the entire period (F_2,211_=0.14, p=0.87). In sharp contrast, the cell density in trained brains (102.4+50, n=65) dropped dramatically after completion of rule learning (F_2,201_=19.4, p<0.001) (Figure 8A_2_, B_1_, B_2_), although it remained significantly higher (t=2.49, p=0.014) than the pre-training value (naïve).

We next asked if the same neurons remain active throughout the learning process, from the onset of training to the completion of rule acquisition. To address this, we double-labeled active neurons at two distinct time points, one in the first day of training using TRAP expression via intraperitoneal injection of 4-hydroxytamoxifen (4-OHT) prior to the training session and the second were subsequently labeled using c-Fos immunohistochemistry (Figure 8C_1_). In this experiment the average number of TRAP2 positive neurons after the first day of training was 196.6±34.5 (32 slices). In the same slices, the average number of c-Fos positive neurons was 114.6±42.7 and the number of double-labeled neurons, positive for both TRAP and c-Fos was 73.2±21.4.

These results show that; ***a.*** Only 37.2%±9.2 of the neurons that were activated at the onset of training are also active in the engram for the rule learning memory (Figure 8C2, left). ***b.*** The 66.5%±11.4 of the neurons that entail the rule learning engram participated throughout the learning process, from the very beginning of training until its completion many days later (Figure 8C2, right). It should be noted that one day after the completion of training, the average number of TRAP labeled neurons (102.4±50.3, n=65) did not differ (t=1.18, p=0.24) from the average number of c-Fos labeled neurons (114.7±42.7, n=32), indicating that the two labeling methods are equaly efficient inlabeling IEGs ecpressing neurons

### IEGs expressing tagged neurons are essential for maintaining the long-term memory of the rule

Finally, we explored the role of IEGs expressing neurons in maintaining the memory of the learned rule by selectively silencing this active neuronal population at the completion of training, using an inhibitory chemogenetic approach (Figure 9A, see methods). An inhibitory DREADD virus was injected into the layer II of the piriform cortex in the TRAP2 mice. After the mice reached the rule-learning criterion. On the day of rule learning completion, 4-hydroxytamoxifen (4-OHT) was injected to induce DREADD expression specifically in neurons activated on the final training day. One day later, clozapine-N-oxide (CNO) was injected to activate DREADD receptors, to silence the tagged neuronal population.

**Figure 9:**
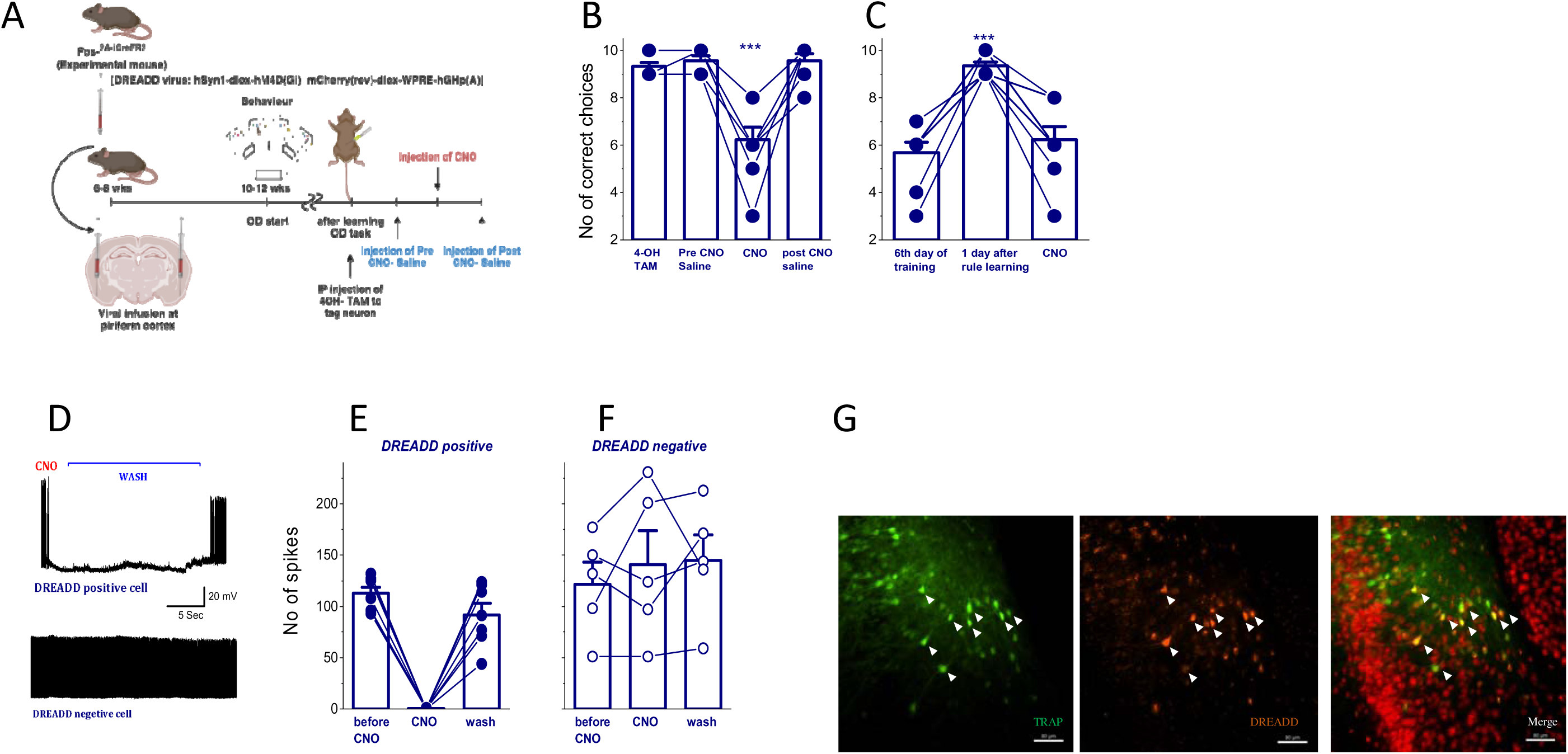
IEGS expressing neurons are crucial for the memory engram of the rule. **A.** The procedure of expressing inhibitory DREADD in the mouse piriform cortex pyramidal neurons (see methods for detail). **B.** DREADD application reduces significantly the level of performance in the OD task after rule learning completion. Each point represents the number of correct choices made by each mouse of the last 10 trials of the session. Bars represent mean ± SE. (*** P<0.001). **C.** For each mouse, DREADD application reversibly reduced the number of correct choices to pre-rule learning performance level. Each point represents the number of correct choices made by each mouse in the last 10 trials of the session. Bars represent mean ± SE. (*** P<0.001). **D**. Examples of the effect of CNO application on repetitive spike firing on a DREADD-positive and a DREADD-negative neuron. **E, F. Quantitative description of the effect of CNO on different neurons.** While CNO application completely shuts down reversibly spike firing in DREADD-positive neurons (E), it has no effect on DREADD-negative neurons (F). Each point represents the number of spikes in each neuron. Bars represent mean ±SE. **G**. Double labeling of TRAP (Green) and DREADD (Red) show the IEGs activated neurons in which spike firing was shut down by DREADD application. Red cells on the right panel were labeled by NeuN (see methods). This shows that the double labeling for TRAP and DREADD is apparent in a small fraction of the cells’ population in the piriform cortex.

First, we examined whether activation of the inhibitory DREADD affects the memory of the learned rule. When CNO was administered two days after rule learning, task performance dropped significantly, well below the level observed in animals that had successfully acquired the rule. Specifically, CNO application reduced significantly the averaged success rate of the task, compared with the performance on the day just before and after CNO (F_3, 29_ =22.2, p=1.4*10^-7^; Figure 9B). In each individual mouse, CNO application reduced the success rate (6.2±1.6, n=9) to the extent recorded on the 6^th^ day during learning (5.7±1.3) (paired t-test, t=0.68, P=0.52; Figure 9C).

These results indicate that targeted silencing the IEGs expressing neurons via inhibitory DREADD significantly impairs the animal’s performance, indicating that this population plays a critical role in memory expression. This effect was reversible; in absence of CNO injections, when inhibitory DREADD were not activated, performance of the animals returned to the typical post rule learning level.

At the cellular level, bath application of CNO (5 µM) completely and reversibly abolished repetitive spike firing, with no effect observed in DREADD negative neuron (Figure 9D-F). Figure 9G shows the specific and localized expression of DREADD positive neuronal cells overlapped with IEGs expressed TRAP-tagged cell, indicating that nearly all IEGs expressing neurons express DREADD.

These findings indicate that the active cell population in the final day of training can be identified as the “neuronal ensemble” for the memory of the rule.

## DISCUSSION

Our findings show that although rule learning requires prolonged and effortful training (Saar et al. 1998;1999; 2001, Chandra and Barkai 2018, Chandra et al. 2019), it appears to emerge almost entirely at a distinct moment marked by the sudden ability to perform the complex tasks in an accelerated learning. Such enhanced ability is supported by long-term changes in all three major components controlling neuronal activity: intrinsic neuronal excitably, synaptic excitation and synaptic inhibition. These changes are expressed in a subset of neurons that form the engram encoding of the long-term memory of the rule.

### Identification of neurons which are central to the memory engram of the rule

Multiple studies have demonstrated the formation of engrams representing context in fear conditioning tasks in the neocortex, hippocampal CA1, and the amygdala (Cai et al., 2016; Ghandour et al., 2019; Grewe et al., 2017; Kitamura et al., 2017). A memory engram is defined as the physical substrate of memory in the brain. It is formed by learning and reactivated by retrieval. During the past decade, it was demonstrated that learning is associated with the induction of immediate-early genes (IEGs), such as cFos and activity-regulated cytoskeleton-associated protein (Arc). The induction of IEGs is tied to encoding (Josselyn et al., 2015) and to subsequent retrieval of the memory engram (Ryan et al., 2015, Meissner-Bernard et al., 2019). Transgenic mice have been engineered where the expression levels of the endogenous IEGs are used as a promoter to drive transcription of a genetically encoded fluorescent protein. This enables neuronal ensembles that were once active to be captured and tagged (Josselyn et al., 2015), allowing to compare between neurons that were active during formation of the memory engram that encodes the behavioral task, and the neighboring neurons that are not part of the engram. Based on the level of IEGs expression, it was found that the subset of cells that participated in the memory engram have enhanced intrinsic excitability (Gouty-Colomer et al., 2016; Benito and Barco, 2010). Several studies showed that freezing behavior in animals trained in a fear conditioning paradigm is correlated with activity of IEGs expressing cells in the lateral amygdala (Han et al., 2009; Kim et al., 2014). These neurons may represent key components in the fear memory traces.

Surprisingly, our data show that IEGs are expressed in more neurons from animals that were exposed to the OD maze and odors regardless if they were trained for the rule (trained as well as pseudo trained); Compared to naïve mice, tagged neurons in brains of trained and pseudo trained mice doubled compared to naive mice. However, the relationship between IEGs expression and the biophysical properties of the neurons in which such experiences were apparent, depends on the context of activation.

For trained animals, IEGs-expressing neurons that are recruited to the learning process, are more intrinsically excitable and show enhanced excitability and reduced AHP from the first days of training throughout the training period, days before the animals show any improvement in their performance. However, for pseudo-trained mice, tagged neurons have the same excitability and AHP as non-tagged neurons. These findings suggest that the exposure to the maze and odors by themselves activated IEGs in piriform cortex neurons. However, only the rule learning process induces reduced AHP, and thus increased excitability. The onset of such r AHP reduction, after the first day of learning when animals are just beginning the long process of learning the rule only, indicates that this intrinsic change does not occur when the animals are exposed to the maze or master the rule.

The process of rule learning possibly involves the secretion of acetylcholine (ACh) onto PC pyramidal neurons, as we have previously shown that the OD rule learning induced enhancement of intrinsic neuronal excitability is mediated by persistent reduction of cholinergic muscarinic current (M-current) (Saar et al. 2001; Awasthi et al. 2022). ACh also has a profound effect on synaptic transmission, excitatory and inhibitory, and synaptic plasticity (Hasselmo 2006; Hasselmo and Stern 2018; Fernández de Sevilla et al. 2021; Kassab 2023). Activation of muscarinic cholinergic receptors induces both long-term potentiation (LTP) and long-term depression (LTD) (Scheiderer et al. 2006; Volk et al. 2007; Fernández de Sevilla et al. 2008; Dickinson et al. 2009; Jo et al. 2010; Sumi and Harada 2023). Notably, the combined effects of ACh have been suggested to support rule learning (Hasselmo and Stern 2018).

Another interesting possibility is that enhanced synaptic input from the olfactory bulb, which is apparent since the beginning of training (Cohen et al. 2015), has a key role in such induction of reduced AHP and enhanced intrinsic excitability (Melyan et al., 2002; 2004; 2011, Chandra et al. 2019). This effect is mediated by metabotropic activation of the GluK2 subtype glutamate receptor (Fisahn et al. 2005; Melyan et al. 2002, 2004, 2011).

### Long-term learning-induced changes in synaptic strength is apparent in IEGs-expressing neurons only

The increase in synaptic strength that mediates memory formation was initially suggested to occur at specific synapses. Notably, learning-induced synaptic modifications include potentiation as well as weakening of synaptic connections (Bear, 1996, Massey and Bashir, 2007), with a relatively small total increase in synaptic excitation onto each neuron. However, a large enhancement of excitatory synaptic strength (>50%) was observed after different training protocols, in several brain areas (McKernan and Shinnick-Gallagher, 1997, Sacchetti et al., 2001; Schroeder and Shinnick-Gallagher, 2005, Cohen et al., 2008,Tye et al., 2008, Yin et al., 2009, Saar et al., 2012, Ghosh et al., 2016, Jammal et al., 2016).

Enhanced synaptic inhibition onto PC pyramidal neurons is also apparent after OD rule learning (Brosh et al. 2009, Saar et al. 2012, Kfir et al. 2014, Ghosh et al., 2015, Reuveni et al. 2013; 2017, Kundu et al. 2024). Such enhanced synaptic transmission is widely spread throughout the pyramidal cell network. However, the strength of inhibition differs significantly between different neurons (Figure 6C, Saar et al. 2012).

Computational analysis raises the possibility that these long-lasting changes appear to be maintained by a whole cell controlled process, rather by a synaptic-specific mechanism (Reuveni et al. 2013; 2017). Such whole cell enhancement of synaptic connectivity is mediated by persistent phosphorylation of the post synaptic AMPA and the GABA_A_ receptors (Ghosh et al. 2015; 2016).

### Rearrangement of the PC network after the completion of rule learning

After rule learning is completed, the sharp step increase in the performance (the “light bulb” moment) is accompanied by a dramatic decrease in the number of IEG expressing neurons in the PC. This network change may be attributed to long-term memory of the rule, not to the fact that the mice are no longer exposed to the maze of the odors, since such reduction does not occur in the brains of the pseudo-trained. Also, such an extent of changes cannot be mediated by a non-specific increase in synaptic inhibition in the cortical circuit; Synaptic inhibition is enhanced onto all pyramidal neurons, tagged and non-tagged, in brains of both trained and pseudo trained mice.

The only synaptic change observed in trained tagged neurons at this time point is an increase in synaptic excitation. Such increase can originate from intrinsic connection between pyramidal neurons (Saar et al. 2001), descending input for the olfactory bulb (Cohen et al. 2015) or ascending connections from the orbitofrontal cortex (OFC) (Wang et al. 2024). Projection to the PC from OFC was shown to have a crucial role in OD learning (Wang et al. 2024). Particularly, the strength from the olfactory bulb is reduced dramatically once the rule is acquired (Cohen et al. 2015). Such a decrease in excitatory synaptic input is likely to reduce the number of active neurons.

The finding that shutting down temporarily the activity of the IEGs expressing neurons only in trained mice erases the memory for the rule, suggests that these neurons compose the memory engram for the rule. Notably, silencing these neurons reduced the performance level to that achieved at the day prior to rule learning (figure 9C). Thus, the decrease in performance is indeed due to temporary blocking of the memory for the rule. It is not the result of a nonspecific effect, such as the depriving the animal from the ability to discriminate between odors or affecting its motivation to perform the task.

### Sequence of events leading to the formation of the long-term memory engram for the rule

It has been suggested that long-term modification in intrinsic excitability and in synaptic strength play important roles in long–term memory induction and maintenance (Lisman et al., 2018, Chandra et al, 2019). Specifically, enhanced intrinsic neuronal excitability enables neuronal ensembles to enter into a state best described as a “*learning mode*” (Chandra et al., 2019), during which activity-dependent synaptic modifications are more likely to occur, as neurons become more readily recruited into emerging memory traces (Yiu et al., 2014; Gouty-Colomer et al., 2016, Chandra et al., 2019).

Our findings support this model. We thus suggest that neurons that are recruited at the very beginning of the extended training period leading to rule learning, immediately undergo a protein synthesis-dependent intrinsic modification (Cohen-Matsliah et al. 2010) resulting in elevated excitability. This early change increases the likelihood that these neurons are reactivated during subsequent training sessions. Consequently, most of the neurons that entail the long-term memory engram for the rule have been recruited from the very beginning of training, allowing the memory engram to emerge gradually and continuously throughout the training period.

For each animal, the realization of the rule occurs at a sharply-defined moment. This realization is accompanied by a significant reduction in the number of activated neurons, most likely due to a transient suppression of synaptic excitation from the olfactory bulb (Cohen et al. 2015). Once the engram is established, the neurons that remain activated receive enhanced excitatory synaptic transmission from the OB, the OFC and intrinsic connections within the PC (Cohen et al., 2008).

On the day immediately following rule acquisition, enhanced synaptic inhibition is observed in all activated neurons, regardless in their involvement in the learning trace (Figure 7B_2_). Over time, however, this enhanced synaptic inhibition becomes restricted to neurons that are part of the engram (Figure 7C_2_).

The strong correlation between synaptic inhibition and excitation within individual neurons suggests that synaptic enhancement, excitatory and inhibitory, arise through a cell-wide mechanism, as previously suggested (Reuveni et al. 2017).

Together, these findings indicate that neurons forming the memory engram are fundamentally different from other pyramidal neurons in the same network. These differences support the stabilization and persistence of the rule-learning memory (Figure 10).

**Figure 10:**
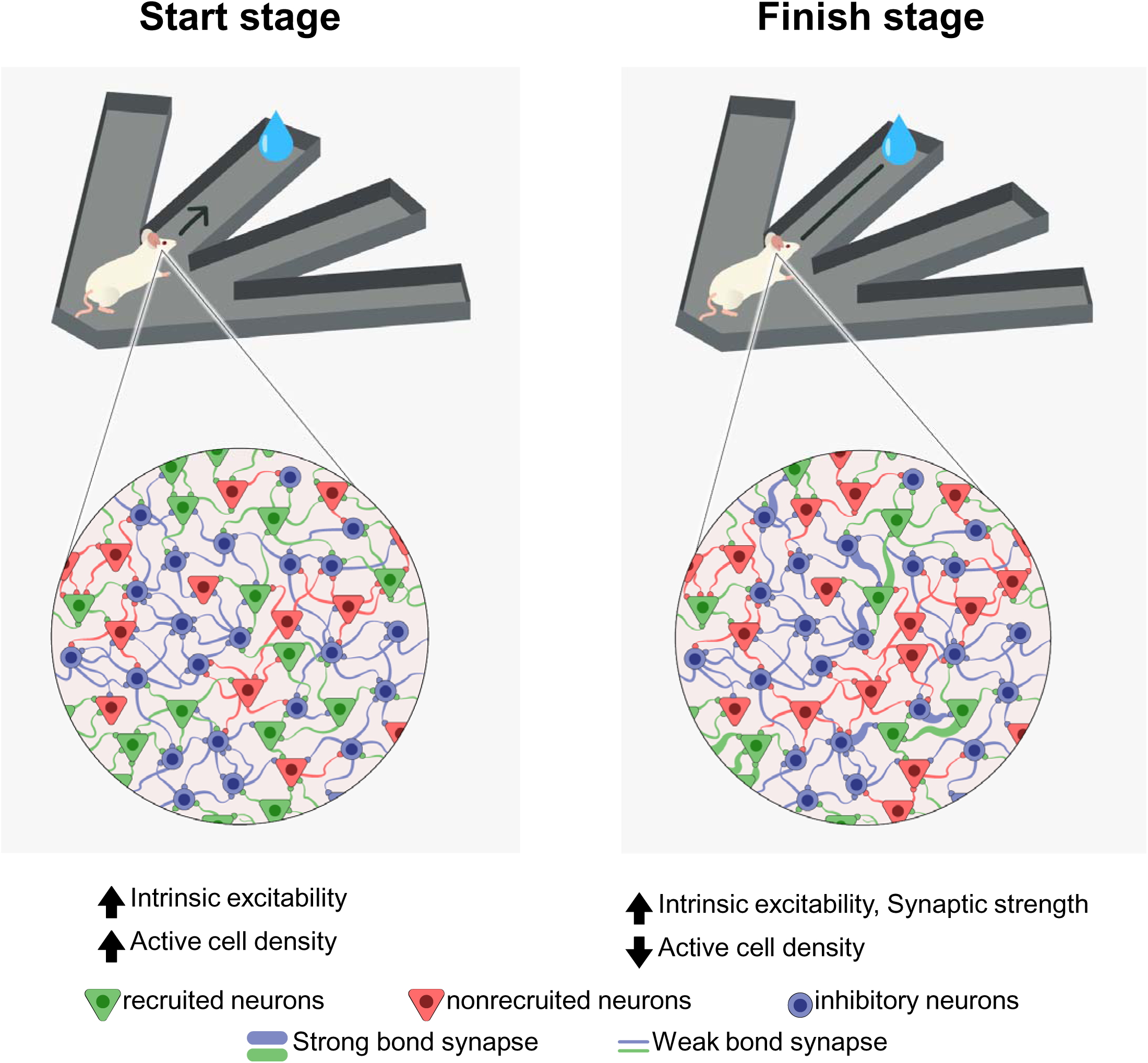
Sequence of cellular and network modifications leading to induction and maintenance of the memory engram of the rule. ***Left:*** From the onset of training, a subset of activated neurons undergrow an intrinsic change, as result of which they become more excitable. This enhanced intrinsic excitably increases the likelihood that these neuros would modify their synaptic connection with neighboring neurons as training for rule learning continues. ***Right:*** Upon completion of rule learning, the long-term memory engram for the rule is composed of a sub group of neurons, most of which were recruited at the onset of training. The number of neurons required to compose the engram is significantly small then the number activated throughout the learning process. The synaptic inputs onto these neurons undergrow a substantial change; The strength of their synaptic connections, excitatory and inhibitory, is enhanced. Notably, excitation and inhibition are increased proportionally in each individual neuron. Such profound change in synaptic connectivity supports the engram stability and thus the long-term memory for the rule.

## EXPERIMENTAL PROCEDURES

### Animals

Hemizygous Fos^2A-iCreER^/ TRAP2 (Targeted Recombination in Active Population) mice strain of both genders were used for all experiments. This mouse line was a gift from Prof. Adi Mizrahi’s laboratory, Hebrew University, Jerusalem, Israel. Mice were group-housed in a cage under temperature and humidity-controlled environment with 12/12 hrs light/dark cycle condition.

Minimum 6-8 weeks aged mice were used (unless mentioned otherwise). Before starting the behavioral experiment, mice were divided into three groups (Trained, Pseudo trained and cage control or Naïve) and were provided with food *ad libitum,* but were maintained on approved water deprivation schedule until the completion of the experiment. All the experimental procedures and animal handlings were as per the guideline by NIH, US and were approved by the University of Haifa animal ethics committee.

TRAP2 mouse line was engineered to have expression of a tamoxifen inducible, improved Cre recombinase (iCre/ERT2) from the “Fos” promoter without disrupting endogenous “Fos” expression. Under the expression of immediate early genes (IEGs; in this case, c-fos) by any specific activities (any sensory or behavioral activity), iCre recombination gene is expressed in a Cre dependent manner discretelyin the presence of tamoxifen, which further allows to express the effector gene of interest, tagged with specific fluorescence molecules to identify the active cell (Guenthner et al., 2013).

### Surgical intervention of stereotaxic virus infusion

To mark the activity dependent neuronal cell and to express the inhibitory DREADD in active neuronal cell, [AAV8/2-hSyn1-dlox-tdTomato(rev)-dlox-WPRE-bGHp(A) (v284, VVF, Zurich), and AAV8/2-hSyn1-dlox-hM4D(Gi)_mCherry(rev)-dlox-WPRE-hGHp(A] (v84, VVF, Zurich) was bilaterally injected respectively, at bregma AP −1.46, ML 3.95, DV 5.20 using a stereotaxic frame (Kopf stereotaxic instrument) (Figure 3B). Further, the expression of labelling active neuronal cells was precisely controlled by IP injection (2mg/kg BW) of 4-hydroxytamoxifen (4 OHT, Sigma-Aldrich; Cat#H6278) (4-OH TAM) to the mouse ∼ 45 mins before starting the behavioral task. Additionally, for inhibitory DREADD activation, CNO (#16882; Cayman) (10 mg/Kg BW) was injected through IP route ∼ 45 mins before starting the behavioral task. Mice were anesthetized under gas anaesthesia machine (SomnoSuite, Kent Scientific, USA) using 4% inhalation anaesthetic (Isoflurane; Piramal Inc.) initially for ∼ 10 mins and then maintained at ∼1.5% throughout surgical procedure. For post-surgery pain management strategy, mice were administered with a weight-adjusted dosage of an anti-inflammatory pain reliever, “Carprofen” (5mg/kg) and antibiotic “Enrofloxacin” (5mg/kg) through IP during the surgery and additional three days post-surgery. The mice were allowed at least 7 days to recover from the surgery and further 3 weeks for expression of viral constructs before any behavioral / electrophysiological / immunohistochemistry experiments were conducted.

### Complex olfactory discrimination (OD) learning and training protocol

OD training was performed in a 4-arm maze (Figure 1A), using commercial synthetic odors that are regularly used in cosmetics and food industries, purchased from Value Fragrance & Flavors. The odours were diluted by ∼420 times with water before applying them in the maze. An OD training protocol was followed similarly to the one mentioned in (Barkai, 1998; Saar et al., 2012) with the following improvisation. Mice 10-12 weeks of age were used for this training task. A day before the training began, mice were provided with a strict 30 mins access to the drinking water and food *ad libitum*, on a daily basis, until the end of the task. The olfactory discrimination training protocol consisted of 20 trials per day. Learning was considered as acquired upon demonstration of at least 80% positive-cue choices for the last 10 trials in a day, referred as the criterion of the task. A pseudo-trained group of age-matched animals were exposed to the same protocol of training, but with random water rewarding. An age-matched naive group of animals were water-deprived but were never exposed to any training protocol. Rule learning required 10 ±2 training days for mice (Figure 1B). Following successful completion of OD training, animals were sacrificed for electrophysiological experiments or for staining experiments.

### In-vitro physiological studies

#### Brain slice preparation and recordings

The overall experimental approach to perform electrophysiological recording starting from the virus injection (activity-induced cell marking) followed by behavioral training and conducting electrophysiological recording, are presented sequentially in figure 3A. Animals were sacrificed under 2% inhalation anaesthetic agent (Isoflurane; Piramal, USA) and the brain slices were prepared. Coronal brain slices of 300 µm thickness for whole cell patch clamp recordings were obtained as previously described (Ghosh et al., 2015; 2016) and were incubated for at least 1 hour prior to recording in oxygenated (95% O2 + 5% CO2) artificial cerebrospinal fluid (aCSF) solution (also referred as bath solution), containing (in mM): 124 NaCl, 3 KCl, 2 MgSO4, 1.25 NaH2PO4, 26 NaHCO3, 2 CaCl2, and 10 Glucose. pH= 7.4, 310 mOsmol. Throughout the recoding period, the bath temperature was maintained at ∼ 30° C using a single channel temperature controller (Warner Instrument Inc.). Recordings were taken from soma of layer-II pyramidal neurons, from every animal from all groups (Figure 3C).

#### Post burst after hyperpolarization potential (AHP) and intrinsic excitability recordings

A single brain slice (one hemisphere) was kept in the recording chamber under infrared DIC fluorescence microscope. The brain slice was perfused with bath solution (∼2ml/min) at a maintained temperature 30°C using single channel temperature controller (Warner Instrument Inc.). Whole cell voltage-clamp recording was done from visually identified soma of pyramidal neuron in layer II piriform cortex (TRAP positive cells – red in color due to presence of tdTom as fluorophore (representative image of marked cell; figure 4A), TRAP negative cells-no colour) by using borosilicate glass (1.5 mm O.D) recoding pipettes (tip resistance 5-6 MΩ) filled with internal solution (Awasthi et al., 2022) (130mM K gluconate, 5mM KCl, 10mM HEPES, 0.6mM EGTA, 2.5mM MgCl2, 4mM Na2ATP, 0.4mM Na3GTP, 10mM sodium phosphocreatine, at pH 7.25, osmolarity ∼290 mOsm, adjusted with KOH) to record post burst AHP and firing frequency.

To record the post burst AHP, cells were held at −60mV in current-clamp mode. With a delay of 200ms, an amount of current was injected to produce 6 spikes within 100ms to burst firing of cell to produce post burst AHP (Figure 4A). A total 10 sweeps were recoded to produce the averaged calculated post burst AHP. The AHP was calculated as the difference in amplitude between the baseline amplitude (before current injection) and the post burst lowest point amplitude. For firing frequency recordings, the cells were held at zero current injection voltage clamp mode. First, the current intensity required to reach the threshold for a single action potential (I_Th_) was determined.

Subsequently, a 1 sec depolarizing pulse was applied, with stimulus intensity of I_Th_X2. Excitability was measured by counting the average number of action potentials evoked by such ten stimuli (Figure 4A). All the recordings were obtained with Axopatch 200B (Molecular Devices) amplifier and data was collected using pClamp 9 (Molecular Devices) software with signal filtered at 5 kHz and sampled at 10 kHz with Axon Digidata 1550B (Axon Instruments).

#### Recording of miniature post-synaptic currents

One hemisphere of a single brain slice was kept in the recording chamber under infrared DIC fluorescence microscope, perfused with bath solution (∼2ml/min) at a maintained temperature of 30°C, using a single channel temperature controller (Warner Instrument Inc.). Whole cell voltage-clamp recording was taken from visually identified soma of pyramidal neuron in layer II piriform cortex (TRAP positive cells – red in colour as mentioned before, TRAP negative cells-no colour) by using borosilicate glass (1.5 mm O.D) recoding pipettes (tip resistance 5-6 MΩ) filled with internal solution (Sakimoto et al., 2019) (127.5mM Cesium methanesulfonate, 7.5mM CsCl, 10mM HEPES, 2.5mM MgCl2, 4mM Na2ATP, 0.4mM Na3GTP, 10mM sodium phosphocreatine, 0.6mM EGTA at pH 7.25, osmolarity 280-285 mOsm) to record the miniature excitatory post-synaptic currents (mEPSC) and miniature inhibitory post-synaptic currents (mIPSC) from the same cell. To record only the mIPSC, 0.5 mM calcium concentration was used instead of 2 mM in aCSF solution. The following recipe was used to prepare the pipette solution; 135 mM CsCl, 1 mM EGTA, 6 mM KCl, 4 mM NaCl, 2 mM MgCl2, and 10 mM HEPES. pH= 7.25. All the recordings were obtained from Axopatch 200B (Molecular Devices) amplifier and data was acquired using pClamp 9 (Molecular Devices) software with signal filtered at 5 kHz and sampled at 10 kHz with Axon Digidata 1550B (Axon Instruments. For both mEPSC and mIPSC recordings, first the mEPSC was recorded for 10 mins in the presence of bath application of TTX (1 µM) voltage clamped the cell at −70 mV. Afterward, the same cell was voltage clamped to 0 mV to record mIPSC for 5 mins in the presence of bath application of TTX (1 µM), DNQX (20 µM) and APV (50 µM) (Saar et al., 2012).

#### Events detection criteria and analysis of the distribution of AHP and IPSCs

For any miniature current recordings, no compensation was adjusted but the series resistance was monitored throughout the experiments in every minute by applying −5 mV 20 ms step current through the recording electrode. Recordings with stable series resistance were considered for analysis. Miniature events were analysed through MiniAnalysis software (formerly-Synaptosoft.com) using automated detection (amplitude, rise and decay time) with visual confirmation of events. Mean amplitude of mEPSC and mIPSC were calculated from each recorded neuron.

To determine the distribution of the AHP amplitudes (Figure 2C) and the IPSCs amplitudes (figure 6C), we have first performed a kernel density estimation using a Gaussian kernel. This yields a densely and evenly sampled pdf (probability density function), which we herewith denote by ***f* (t)**. Since, for all data sets, we seek a bimodal normal distribution, ***f* (t)** assumes the form

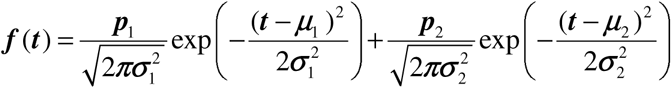

Where, for ***i*** =1,2, ***p_i_*** is the weight of the corresponding unimodal Normal distribution (i.e the ratio of the data obeying it), and σ***_i,_ µ_i_*** its standard deviation and expectation.

We have opted to directly fit the model to the discrete pdf, by minimizing the fit error defined by 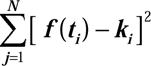, where ***t****_i_*, ***k****_i_* are the data values and associated probabilities produced by the kernel density estimation (we used ***N*** = 512). In order to reach a global minima, we ran many starting points for the minimization process, which cover a five-dimensional grid (note that ***p***_1_ + ***p***_2_ = 1, hence there are five independent variables to minimize over). Then, we ran a gradient descent-based algorithm starting at each grid point and chose the parameters with the minimal function value. To exclude the possibility of missing the global minimum, we have bounded from below the value at each grid box, by using the multivariable version of the mean value theorem.

We note that while highly accurate, the computational complexity of the described method increases exponentially with the number of the parameters.

### Immunohistochemistry (IHC) protocol

#### cFOS staining

IHC was performed against cFos protein for its detection to perform the co-localization study with activity dependent marked neuronal cell. Mice were sacrificed 90 mins after the behavioral performance of the OD maze (Figure 8) and were intracardially perfused with cold phosphate-buffered saline (PBS) solution followed by cold 4% paraformaldehyde (PFA) dissolved in PBS. The brains were removed and post-fixed in a 4% PFA solution overnight at 4°C. Following post fixation, brains were sliced on a vibrotome into 40-μm-thick sections. Every third section was collected in a 24 well-plate (Biofil) and the free-floating brain sections were rinsed in PBS solution for 3 times, 10 mins each. Following rinsing, sections were pre-incubated in a blocking solution (20% goat serum albumin; Biological Industries and 0.3% Triton X-100; Fluka in PBS) for 2 hrs in room temperature. In the post blocking procedure, sections were treated with an anti-c-Fos antibody (Cell Signaling Tech, 9F6, Cat. #2250, 1:500) in blocking solution overnight at 4°C in a shaker. The sections were rewashed in PBS solution for 3 times, 10 mins each and again incubated with goat anti-rabbit IgG antibody coupled with Alexa Fluor 488 (ab150081, Abcam. 1:1000) in PBS for 1 hr at room temperature. Following treatment with secondary antibody, slices were mounted on clean glass slides, air-dried, and cover slipped with aqueous mounting medium (Fluoromount #F4680; Sigma) mixed with 2µM Hoechst 33342 Solution (Cat# 62249; Thermo Fisher) for imaging. Images of the whole sections were obtained under 10X magnification using Nikon Eclipse Ti2-E inverted wide field fluorescence microscope.

#### NeuN staining

Immunofluorescence staining was performed against neuronal nuclei antigen (NeuN) to quantify neurons in the layer II-III in the piriform cortex of both hemispheres of the mouse brain. Mice were sacrificed after the completion of the OD learning task and were intracardially perfused with cold PBS solution followed by cold 4% PFA dissolved in PBS. The brains were removed and post-fixed in a 4% PFA solution overnight at 4°C. Following post fixation, brains were sliced on a vibrotome into 40-μm-thick sections. Every third sections were collected in a 24 well plate (Biofil) and the free-floating brain sections were rinsed in PBS solution 3 times, for 10 mins each. Following rinsing, sectioned were pre-incubated in a blocking solution (20% normal goat serum; Biological Industries and 0.3% Triton X-100 in PBS) for 2 hrs in room temperature.

Post blocking, sections were treated with anti-NeuN primary antibody (abcam, Cat. # Ab177487, 1:500) in blocking solution overnight at 4°C in a shaker. The sections were rewashed in PBS solution for 3 times, 10 mins each and again incubated with goat anti-rabbit IgG antibody coupled with Alexa Fluor 647 (ab150079, Abcam. 1:1000) in PBS for 2 hr at room temperature. Following treatment with secondary antibody, slices were rewashed in PBS solution for 3 times and then mounted on clean glass slides, air-dried, and cover slipped with the mounting medium (Fluoroshield™ with DAPI #F6047; Merck) for imaging

### Statistical analysis

All the data were analysed in MS office 365 excel file and statistic was performed using OriginLab software. ‘n’ refers to the number of cells, unless mentioned otherwise. 2-3 cells were recorded from each animal. One-way ANOVA was used to evaluate the significance of difference between three populations (i.e. trained, pseudo-trained and naive groups), following a normality test. When the ANOVA test indicated that significant differences exist, post hoc Tukey’s test was performed to determine if there’s a difference between each pair of the three groups. Difference between groups where data were non-normally distributed was determined using the Kolmogorov-Smirnov test. Learning-related modifications were considered to occur when the trained group differed from the pseudo-trained group and the naive group and the two control groups were not different. Data presented in the text mentioned as “mean ±SD” and in the figures as “mean ±SE” In all comparisons, the significance level was defined as p value less than 0.05and abbreviated as: *P <0.05, **P <0.01, ***P <0.001.

## Notes

### Competing Interest Statement

The authors have declared no competing interest.

### Funding

- Israel Science Foundation

